# Transcriptomic analysis of native versus cultured human and mouse dorsal root ganglia focused on pharmacological targets

**DOI:** 10.1101/766865

**Authors:** Andi Wangzhou, Lisa A. McIlvried, Candler Paige, Paulino Barragan-Iglesias, Carolyn A. Guzman, Gregory Dussor, Pradipta R. Ray, Robert W. Gereau, Theodore J. Price

**Affiliations:** The University of Texas at Dallas, School of Behavioral and Brain Sciences and Center for Advanced Pain Studies, 800 W Campbell Rd. Richardson, TX, 75080, USA; Washington University Pain Center and Department of Anesthesiology, Washington University School of Medicine

## Abstract

Dorsal root ganglion (DRG) neurons detect sensory inputs and are crucial for pain processing. They are often studied *in vitro* as dissociated cell cultures with the assumption that this reasonably represents *in vivo* conditions. However, to our knowledge, no study has ever directly compared genome-wide transcriptomes of DRG tissue *in vivo* versus *in vitro*, or between different labs and culturing protocols. We extracted bilateral lumbar DRG from C57BL6/J mice and human organ donors, and acutely froze one side and processed the other side as a dissociated cell culture, which was then maintained *in vitro* for 4 days. RNA was extracted and sequenced using the NextSeq Illumina platform. Comparing native to cultured human or mouse DRG, we found that the overall expression level of many ion channels and GPCRs specifically expressed in neurons is markedly lower in culture, but still expressed. This suggests that most pharmacological targets expressed *in vivo* are present in culture conditions. However, there are changes in expression levels for these genes. The reduced relative expression for neuronal genes in human DRG cultures is likely accounted for by increased expression of genes in fibroblast-like and other proliferating cells, consistent with the mitotic status of many cells in these cultures. We did find a subset of genes that are typically neuronally expressed, increased in human and mouse DRG cultures, including genes associated with nerve injury and/or inflammation in preclinical models such as *BDNF*, *MMP9*, *GAL,* and *ATF3*. We also found a striking upregulation of a number of inflammation-associated genes in DRG cultures, although many were different between mouse and human. Our findings suggest an injury-like phenotype in DRG cultures that has important implications for the use of this model system for pain drug discovery.

## Introduction

Nociceptors within the dorsal root ganglia (DRG) or trigeminal ganglia (TG) are the first neurons in the pain pathway and are responsible for the detection of damaging and potentially damaging stimuli [17; 62]. These neurons are crucial contributors to chronic pain disorders ranging from inflammatory to neuropathic pain [46; 48]. Because of their importance in clinical pain disorders, these neurons are frequently studied to gain insight into mechanisms that drive chronic pain and to develop better treatment strategies. Traditionally, investigators have studied rodent nociceptors *in vitro* as dissociated cell cultures prepared from DRG or TG. More recently, investigators have also started to study DRG nociceptors from human organ donors and surgical patients [14; 39; 47; 50; 51; 59; 70]. This creates a “clinical bridge” for advancing mechanisms or therapeutics from rodents toward the clinic. These models have many advantages; cultures can easily be used for electrophysiology, Ca^2+^ imaging, biochemical, or other functional studies. These studies have unquestionably advanced the field of pain neurobiology and sensory transduction.

Despite the widespread use of this model system [35], many investigators are skeptical of the degree to which these cells in dissociated culture accurately reflect the status of nociceptors *in vivo*. Several studies have analyzed the genome wide RNA profiles of these cultures [25; 43], but not in the context of changes with respect to the intact ganglia. A previous study by Thakur *et al* [54] contrasted RNA sequencing (RNA-seq) profiles of intact DRGs with unsorted, dissociated DRGs in the context of profiling magnetically sorted, neuronally enriched dissociated DRGs. The study found few differences (7,630 genes were comparably expressed in both, while 424 were differentially expressed) between intact DRG tissue and unsorted, acutely dissociated DRG, suggesting that the process of dissociation does not dramatically alter the molecular phenotype. While some studies have compared expression of a single gene or a handful of genes in these *in vitro* cultures vs. the intact ganglia (like the comparison of *Npr3* expression in Goswami *et al* [22]), we are unaware of any study that has used genome-wide assays to study how gene expression might be altered from native to cultured DRG conditions. In this study we addressed this question by comparing intact versus cultured DRG from human donors and mice using RNA-seq technology. We designed a series of experiments to study how native DRG transcriptomes differ from cultured ones in humans and mice. Our findings provide a comprehensive, genome-wide evaluation of gene expression changes from native to cultured DRG in both humans and mice. Consistent with previous studies [18; 40], we found that DRG neurons in culture show transcriptional signatures that suggest a neuropathic pain phenotype [5; 26]. This supports the use of cultured DRG neurons as a model system to study underlying mechanisms of pain. However, our findings point out some shortcomings of using these models to study multiple classes of receptors that show altered expression in culture. Some of these differences do not occur consistently across species, suggesting mouse DRG cultures may not be a good surrogate for human cultures in certain experiments. The data provided in this study is presented in a companion website (https://bbs.utdallas.edu/painneurosciencelab/sensoryomics/culturetxome/) that will help investigators choose and design appropriate experimental parameters, and can provide an important tool for future experiments in pain and somatosensory fields.

## Methods

### Experimental Design

Because genetic variation can be a possible contributor to transcriptome level differences in nervous system samples from human populations [39; 41], we chose a study design wherein we cultured lumbar DRGs from one side in human donors and immediately froze the opposite side from the same donor for RNA sequencing. Although we used an inbred mouse strain (C57BL/6) for parallel mouse studies, we used a similar culturing design where cultures were done in two independent laboratories to look for variability across labs. RNA sequencing was performed at 4 days in vitro (DIV) to stay within the electrophysiologically relevant range of 1 – 7 DIV for human DRG and the biochemical assay range of 4 – 7 DIV for both human and mouse DRG.

### Animals

Price Lab: All procedures were approved by the Institutional Animal Care and Use Committee of University of Texas at Dallas and were in strict accordance with the US *National Institute of Health (NIH) Guide for the Care and Use of Laboratory Animals*. Adult C57Bl/6 mice (8-15 weeks of age) were bred in house, and were originally obtained from The Jackson Laboratory. Animals were housed in the University of Texas at Dallas animal facilities on a 12 hour light/dark cycle with access *ad libitum* to food and water.

Gereau Lab: All procedures were approved by the Animal Care and Use Committee of Washington University and in strict accordance with the US *National Institute of Health (NIH) Guide for the Care and Use of Laboratory Animals*. Adult C57Bl/6 mice (8-15 weeks of age) were bred in house, originally obtained from The Jackson Laboratory. Animals were housed in Washington University School of Medicine animal facilities on a 12 hour light/dark cycle with access *ad libitum* to food and water.

### Intact vs cultured mouse DRG

Price lab: Male and female C57BL/6 mice (4 week-old, ∼15-20 g; n=3, for each sex) were anesthetized with isoflurane and killed by decapitation. Lumbar DRGs (L1-L6) from one side of the spine were frozen in RNAlater (Invitrogen) while DRGs from the other side from the same mouse was cultured and then scraped at 4 DIV into RNAlater. L1-L6 DRGs for culturing were dissected and placed in chilled HBSS (Invitrogen) until processed. DRGs were then digested in 1 mg/ml collagenase A (Roche) for 25 min at 37°C then subsequently digested in 1 mg/ml collagenase D for 20 min at 37°C. DRGs were then triturated in 1 mg/ml trypsin inhibitor (Roche), then filtered through a 70 μm cell strainer (Corning). Cells were pelleted then resuspended in DMEM/F12 with GlutaMAX (Thermo Fisher Scientific) containing 10% fetal bovine serum (FBS; Thermo Fisher Scientific), 1% penicillin and streptomycin, 5 ng/mL mouse 2.5S NGF (Millipore), and 3 μg/ml 5-fluorouridine with 7 μg/ml uridine. Cells were distributed evenly across 4 wells using a 24-well plate coated with poly-D-lysine (Becton Dickinson). DRG neurons were maintained in a 37°C incubator containing 5% CO2 with a media change every other day. At 4 DIV, cells were scraped into 500 uL RNAlater and processed for RNA extraction.

Gereau lab: Male and female C57Bl/6 mice (n=3, for each sex) were deeply anesthetized with isoflurane and quickly decapitated. From one side, L1-6 DRG were extracted, directly placed into 500μL RNAlater, and stored at −80°C. From the other side, L1-6 DRG were extracted and dissociated in freshly made *N*-methyl-D-glucamine (NMDG) solution (Valtcheva et al 2016). DRG were digested in 15U/mL papain (Worthington Biochemical) for 20min at 37°C, washed, and then further digested in 1.5 mg/mL collagenase type 2 (Sigma) for another 20 min at 37°C. DRG were washed and triturated in DRG media [5% fetal bovine serum (Gibco) and 1% penicillin/streptomycin (Corning) in Neurobasal A medium 1x (Gibco) plus Glutamax (Life Technologies) and B27 (Gibco)]. Final solutions of cells were filtered (40 µm, Fisher) and cultured in DRG media on coverslips coated with poly-D-lysine (Sigma) and rat tail collagen (Sigma). Cultures were maintained in an incubator at 37°C containing 5% CO_2_. On 4th DIV (no media changes), cultured coverslips were scraped in 500 μL RNA later and stored at −80°C.

### Intact vs cultured human DRG

Studies involving human DRG were done on de-identified biospecimens and approved by Institutional Review Boards at Washington University in St. Louis and University of Texas at Dallas.

Gereau lab: Human dorsal root ganglia extraction and culturing was performed as described previously (Valtcheva et al 2016), in a similar manner to the mouse culturing protocol. Briefly, in collaboration with Mid-America Transplant Services, L4-L5 DRG were extracted from tissue/organ donors less than 2 hrs after aortic cross clamp. Donor information is presented in Table 1. DRGs were placed in NMDG solution for transport to the lab for fine dissection. From one side, intact L4-5 DRG were directly placed into 500 μL RNAlater, and stored at −80°C. From the other side, L4-5 DRG were minced and cultured. Pieces were dissociated enzymatically with papain and collagenase type 2 for 1hr each, and mechanically with trituration. Final solutions were filtered (100 µm, Fisher) and cultured with DRG media. On DIV4, cultured coverslips were scraped in 500μL RNAlater and stored at −80°C.

**Table 1.** Human DRG donor characteristics and donor – sample mapping

| Donor id | Age | Sex | Race | Cause of Death | Sample ids |
| --- | --- | --- | --- | --- | --- |
| 1 | 53 | F | White | ICH/Stroke | hDRG-1F, hDRG-1Fre, hDIV4-1F, hDIV4-1Fre |
| 2 | 12 | F | White | Anoxia/OD | hDRG-2F, hDIV4-2F |
| 3 | 26 | M | White | Head trauma/MVA | hDRG-3M, hDIV4-3M |
| 4 | 34 | M | White | Anoxia/OD | hDRG-4M, hDIV4-4M |
| 5 | 18 | F | White | Head trauma/MVA | hDRG-5F, hDIV4-5F |
| 6 | 18 | M | White | Head trauma/GSW | hDRG-6M, hDIV4-6M |

### RNA sequencing

Human and mouse DRG tissue/cultured cells were stored in RNAlater and frozen in −80 °C until use. Samples obtained at the Washington University at St Louis were shipped to UT Dallas on dry ice for uniform library preparation. All RNA isolation and sequencing was done in the Price Lab. On the day of use, the frozen tubes were thawed to room temperature. To obtain RNA from tissue samples, the tissue was extracted from RNAlater with ethanol cleaned tweezers and put in 1 mL of QIAzol (QIAGEN Inc.) inside 2 mL tissue homogenizing CKMix tubes (Bertin Instruments). To obtain RNA from cell cultures, cells were spun down to the bottom of the tube by centrifuge at 5000 x g for 10 min. RNAlater was then removed from the tube, and cells were resuspended with 1 mL of QIAzol and transferred to the homogenizing tube. For both tissues and cell cultures, homogenization was performed for 3 x 1 min with Minilys personal homogenizer (Bertin Instruments) at 4 °C. This time course was used to avoid heating during homogenization. RNA extraction was performed with RNeasy Plus Universal Mini Kit (QIAGEN Inc.) with the manufacturer provided protocol. RNA was eluted with 30 µL of RNase free water. Based on the RNA size profile determined by the Fragment Analyzer (Agilent Technologies) with the High Sensitivity Next Generation Sequencing (NGS) fragment analysis kit, we decided to sequence all human samples with total RNA library preparation and all mouse samples with mRNA library preparation. Total RNA was purified and subjected to TruSeq stranded mRNA library preparation for mouse or total RNA Gold library preparation (with ribosomal RNA depletion) for human, according to the manufacturer’s instructions (Illumina). Quality control was performed for RNA extraction and cDNA library preparation steps with Qubit (Invitrogen) and High Sensitivity NGS fragment analysis kit on the Fragment Analyzer (Agilent Technologies). After standardizing the amount of cDNA per sample, the libraries were sequenced on an Illumina NextSeq500 sequencing platform with 75-bp single-end reads in multiplexed sequencing experiments, yielding at least 20 million reads per sample. mRNA library preparation and sequencing was done at the Genome Center in the University of Texas at Dallas Research Core Facilities.

### Computational analysis

#### Mapping and TPM quantification

RNA-seq read files (fastq files) were checked for quality by FastQC (Babraham Bioinformatics, https://www.bioinformatics.babraham.ac.uk/projects/fastqc/) and read trimming was done based on the Phred score and per-base sequence content (base pairs 13 through 72 were retained). Trimmed Reads were then mapped against the reference genome and transcriptome (Gencode vM16 and GRCm38.p5 for mouse, Gencode v27 and GRCh38.p10 for human [21]) using STAR v2.2.1 [16]. Relative abundances in Transcripts Per Million (TPM) for every gene of every sample was quantified by stringtie v1.3.5 [44]. Downstream analyses were restricted to protein coding genes to make human (total RNA) and mouse (polyA+ RNA) libraries comparable, hence TPMs of only genes annotated as coding genes in the Gencode database were renormalized to sum to a million.

#### Hierarchical clustering

RNA-seq samples for each species were analyzed for similarity by performing hierarchical clustering. The distance metric used for clustering was (1 – Correlation Coefficient) based on Pearson’s Correlation Coefficient [42], and average linkage was used to generate the dendrogram from the distance matrix.

#### Outlier analysis

In human cultured DRG samples, we detected an outlier (sample id hDIV4-1F, Fig. 1A). To rule out incorrect library construction, we sequenced this sample again using another independently prepared library. However, the new library was still an outlier upon sequencing but very similar to the original library (suggesting low technical variability in our library preparation and sequencing steps). In contrast to the other human DRG cultures, this sample had negligible expression levels for many neuronal markers like *CALCA, TRPV1,* and *SCN10A* (**Supplementary file 1, sheet 1)** suggesting that few neurons survived the culturing process for this sample. Consistent with this, experimental notes regarding cultures from hDIV4-1F indicated very sparse apparent neurons in the cultures (not shown). Thus, this sample and its paired acutely dissected sample (sample id hDRG-1F), were excluded from further analysis. A mouse outlier sample (sample id mDIV4-4Fg, Fig. 1 B) was similarly analyzed, but expression of neuronal marker genes was considered sufficient for retention in the analysis.

**Figure 1.**
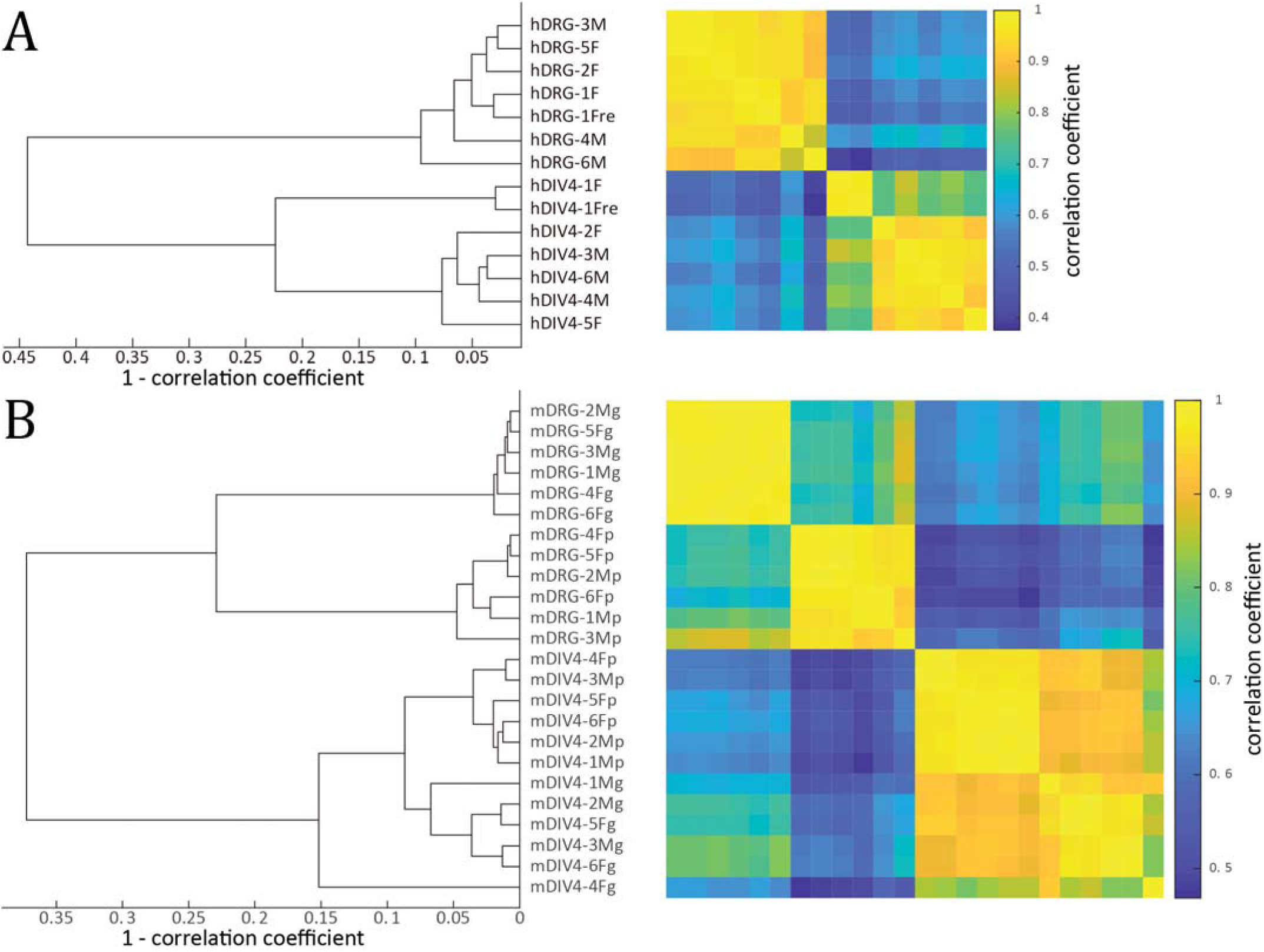
Hierarchical clustering of all human (**A**) and mouse (**B**) samples based on TPM-based whole genome gene abundances. **A.** Cultured and acutely dissected human DRG tissue samples are separated into two clusters. The outlier sample hDIV-1F and its paired dissected sample (hDRG-1F) were excluded from further analysis. **B.** Cultured and acutely dissected mouse DRG samples also segregate into separate clusters. Subclusters in the cultured DRG and dissected DRG clusters correspond to sample generated in Gereau and Price laboratories. The outlier sample mDIV4-4Fg shows moderate expression of neuronal genes, and clusters with other Gereau laboratory cultured samples when unrooted clustering is performed for cultured mouse DRG samples. (Sample id nomenclature -- Prefix: h - human; m - mouse; Infix: DRG - acutely dissected DRG samples; DIV4 - 4 days*in vitro* DRG cultures; Suffix: M - male; F - female; p - Price laboratory; g - Gereau laboratory; re – repeated library preparation and sequencing)

#### Identification of consistently detectable genes

Previous studies on whole DRG tissue have found functional responses for GPCRs with < 0.4 TPMs (e.g. *GRM2* functionally studied and abundance quantified in the papers [14; 47]). This suggests that the approach of picking an expression threshold (in TPMs) to classify a gene either as “on” or “off” is likely to miss functionally relevant gene products based on traditional thresholds (∼ 1 TPM, as in North *et al* [39]). Instead, we classified consistently detectable genes based on reads being detected in the exonic region in 80% or more of the samples in a particular condition (i.e. in at least 4 of 5 human replicates, or in at least 10 of 12 mouse replicates). Assuming iid probabilities for detecting a read emanating from a particular gene in an RNA-seq experiment, this criterion causes the sensitivity of our approach to be suitable for our purpose, calling consistently detectable genes to be those that have ≥1 read in 7 million coding gene reads in an RNAseq library, as :

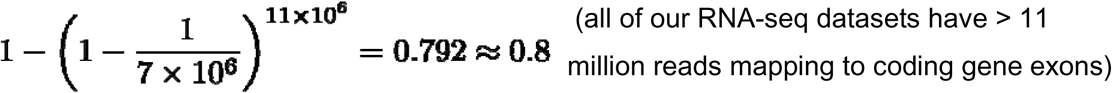

#### Differential expression metrics

Due to small sample sizes in humans, stringent statistical hypothesis testing using Student’s t test [53] with Benjamini-Hochberg multi-testing correction [3] yield few statistically significant differences.

We therefore decided to use strictly standardized mean difference (SSMD) to discover genes with systematically altered expression levels between experimental conditions. For each human and mouse coding gene, we report fold change and the SSMD across conditions. SSMD is the difference of means controlled by the variance of the sample measurements. We used SSMD as a secondary effect size since it is well suited for small sample sizes as in our human samples [39; 71], while simultaneously taking into account the dispersion of the data points. For determining SSMD thresholds that identify genes that are systematically changing between conditions, we use the notion of the related Bhattacharyya coefficient [6], which is used to calculate the amount of overlap in the area under the curve of the two sample distributions in order to control for false positives in differential expression analysis. For homoskedastic Gaussian distributions, we find that based on the Bhattacharyya coefficient, the less stringent constraint | SSMD | > 2.0 corresponds to a 36.8% overlap in the area under the curve of the two sample distributions being tested, while the more stringent | SSMD | > 3.0 corresponds to a 10.5% overlap. The less stringent criterion was used to select differentially expressed genes in gene sets of pharmacological interest, since genes with a moderate amount (< 36.8%) of overlap in TPM distributions between acutely dissociated and cultured DRG should likely not be targeted for pharmacological purposes. The more stringent constraint corresponding to little or no overlap in sample distributions (<10.5%) was used to identify differentially expressed genes at the genome wide level.

Since our data is paired, we report several variations of the standard fold change metric. We calculated the ratio of means across conditions to compare cohort level statistics, but also calculate the mean of ratios of paired samples to better control for individual to individual variations in the transcriptome. However, the mean of ratios is more susceptible to outlier values, so we further modified it to calculate the median of ratios. All fold changes are reported as log_2_ fold changes, for symmetric scaling of fold changes in both directions. Since naïve filtering or ranking by log-fold change can produce incorrect results [45], we constrain differentially expressed genes by SSMD threshold. However, we do additionally constrain that the fold change (ratio of means or median of ratios) be > 1.5, since dosage-based functional effects are unlikely to be manifested as a result of lower fold changes.

To avoid issues in calculations of these metrics for genes with no detectable reads in one or both conditions, a smoothing factor of 0.01 was added to both the numerator and denominator when calculating fold changes, and to the denominator when calculating the SSMD. We also provide uncorrected *p* values for paired, two sample, two tailed t tests conducted for individual genes.

These cohort and inter-cohort statistics, along with individual sample TPMs, and cohort means, are provided in **Supplementary file 1, sheets 1 and 2**.

#### Estimation of density functions

To estimate the density functions of fold change (ratio of means) and SSMD for human and mouse pharmacologically relevant genes, we used the inbuilt ksdensity function in Matlab, using normal kernel smoothing.

#### Human – mouse gene orthology mapping and gene expression change comparisons across species

Orthologous genes with a one-to-one mapping between human and mouse genomes were identified using the Ensembl database [24]. Genes from the relevant gene families (GPCRs, ion channels, kinases) were removed from analysis if one-to-one orthology was not identified between human and mouse genes. Additionally, due to the complicated nature of the orthology map in the olfactory receptor and TAS2R families in mice and human [13; 67], these genes families were also excluded from analysis. For all remaining genes in these families that were consistently detected in human or mouse samples, a trend score was calculated by multiplying the SSMD and log median of paired fold change values. The correlation of the human and mouse trend scores were calculated using Pearson’s R [42]. Genes not systematically detected in samples of either species were left out of the analysis to avoid inflating the correlation based on the trend scores.

#### Gene list compilation

Gene lists used in the paper (ion channel, GPCR, kinase) were acquired from online databases including the Gene Ontology (AMIGO), HUGO Gene Nomenclature Committee (HGNC) and the Human Kinome database [19; 32; 55].

Marker gene lists for constituent cell types in the DRG were sourced from the literature and validated in a recently published mouse nervous system single cell RNA-seq database [69]. CNS microglia marker genes were used as a surrogate for marker genes in PNS macrophages (not profiled in the database). We found that many of the traditional protein-based fluorescence markers for these cell types were not ideal for our analyses. Based on our own analysis in the Zeisel *et al* database, the literature-based markers *Gap43*, *Ncam1* and *Ncam2* for non-myelinating Schwann cells were also found to be expressed in Satellite Glial Cells (SGCs) and/or neurons, and were not used. Similarly, SGC markers *Dhh*, *Fbln5*, and *Ceacam10* are expressed in both Schwann cells and SGCs and not used. *Fbnl2*, *Tyrp1*, and *Prss35* were found to be comparably enriched in proliferating and non-proliferating SGCs. Microglial / macrophage markers *Apoe*, *Fabp7* and *Dbi* were also found to be expressed in SGCs and not used as markers. Finally, *Trpc5* was found to be absent in mouse sensory neurons.

Coding was done in Matlab, and data visualization was performed in Matlab and GraphPad Prism V8.

## Results

### Hierarchical clustering of human and mouse samples reveal whole transcriptome differences between cultured and acutely dissected DRG

We used hierarchical clustering to assess differences between RNA-seq samples analyzed in this study. As shown in Figure 1, the top-level split of the hierarchical clustering for both human and mouse samples was between cultured and acutely dissected DRG tissue, showing consistent whole transcriptome changes between the two. We identified broad changes in the transcriptome between acutely dissected and cultured DRGs, with 2440 human and 2941 mouse genes having a fold change (ratio of means and median of ratios) > 1.5, and | SSMD | > 3.0 between compared conditions (**Supplementary file 1, sheets 1 and 2**). The smaller number of changed genes that we detect in human can be attributed to a smaller number of detected genes that increase in abundance in culture in humans compared to mouse. Of the differentially expressed genes, only 443 (18%) of the human genes and 1156 (39%) of the mouse genes have increased abundances in cultured conditions, which suggests that a majority of the differentially expressed genes gain in relative abundance in acutely dissected DRGs compared to culture. Controlled laboratory conditions and a similar genome (belonging to the same mouse strain) potentially causes lower within-group variation at the level of individual genes in the mouse samples with respect to the human samples. The smaller number of human genes detected to be increasing in culture can likely be attributed to higher within-group variation in human samples, since genes that show significantly increased expression in cultured conditions have more moderate changes (median across ratio of means in genes satisfying differential expression criterion - human: 2.8 fold, mouse: 3.5 fold) in expression compared to genes that show significantly increased expression in acutely dissected DRGs (median in human: 5.4 fold, mouse: 5.1 fold). They are therefore less likely to be detected in a lower signal to noise ratio scenario.

### No distinct differences at the whole transcriptome level across sexes

In both human and mouse samples, we did not find clear sex differences at the whole transcriptome level though individual sex markers like *UTY* differ between the sexes (**Supplementary file 1, sheets 1 and 2**), consistent with previous findings [31]. Thus, male and female samples were grouped together for further analyses.

### Increases in SGC and fibroblast markers compensated for by decrease in neuronal and Schwann cell markers in human and mouse cultures

Due to the magnitude of changes, we tested whether the proportion of mRNA sourced from the different constituent cell types of the DRG were different between acutely dissected and cultured samples. We profiled the expression levels of neuronal, fibroblast, Schwann cell, SGC, and macrophage marker gene panels (chosen based on mouse single cell profiles [69]) in both human and mouse cultured and acutely dissected DRGs. We found that neuronal markers were broadly downregulated in all cultured samples from mice and humans. Expression levels of human neuronal markers in culture were decreased by a median of 8.16 fold (Fig 2A). Conversely, markers for human fibroblast-like cells (often of vascular origin) were increased by a median of 4.18 fold (Fig 2B) in culture compared to acutely dissected samples. We found that human myelinating Schwann cell markers (*MPZ*, *MBP*) in culture were decreased by a median of 9.88 fold compared to intact tissues (Fig 2C) but markers for human SGCs, especially proliferating SGCs (Fig 2D), were increased (by a median of 11.60 fold). These trends were conserved in the mouse datasets as well. Since these changes happen broadly (as shown by the density function across pharmacologically relevant gene families, Fig 3) and not just in specific regulatory pathways or gene sets, they indicate that the proportion of mRNA derived from neurons (and possibly Schwann cells) in our RNA-seq libraries decreases in cultured samples. In turn, this suggests that the proportion of neurons (which are post-mitotic) to other cell types decreased in DRG cultures, while the proportion of dividing cells (such as fibroblast-like cells and SGCs) to other cell types increased. However, Schwann cells, which can be mitotic and proliferate under certain conditions, potentially also decrease in proportion based on our data. This is likely because axonal contact is required for Schwann cell survival [64]. Developmentally established transcription factor expression that define sensory neuronal identity (*PRDM12*, *TLX2*, *TLX3*, *POU4F1*, *DRGX*) are all consistently decreased in human and mouse cultures (**Supplementary File 1 Sheets 1 and 2**), further suggesting that the observed changes are more likely to be caused by changes in relative proportions of cell types rather than molecular plasticity of neurons. These changes were expected, given the different mitotic statuses of these cell types, and were almost certainly the primary factors in distorting the transcriptome from what is seen *in vivo*. The zero-sum nature of our relative abundance measure (transcripts per million) potentially also amplifies this signal. Changes in macrophage markers are discussed in a following subsection discussing the pro-inflammatory phenotype of cultured DRGs.

**Figure 2.**
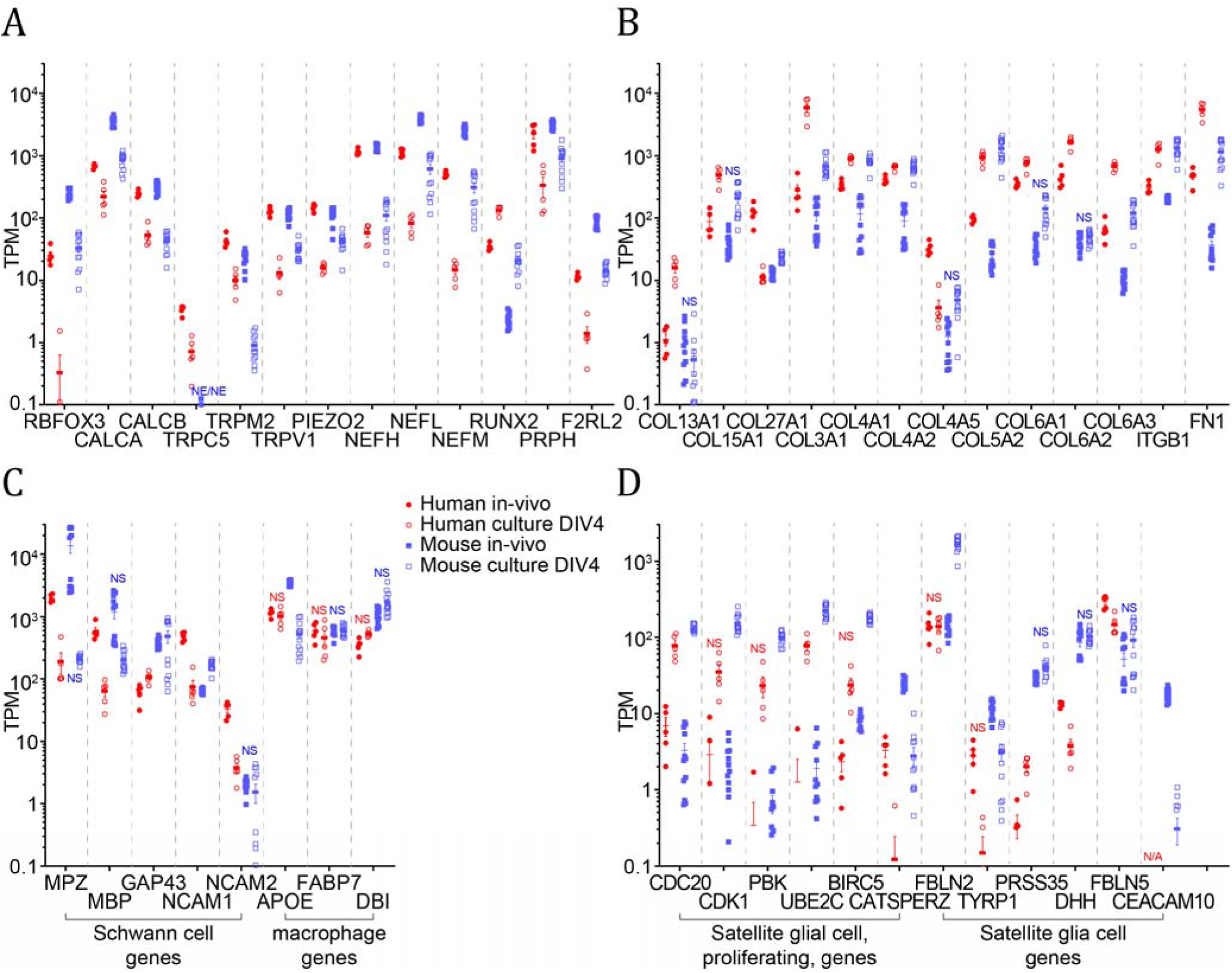
Marker gene expression in dissected and cultured human and mouse DRGs, for neurons (**A**), fibroblast-like vascular cells (**B**), Schwann cells and microglia (as a surrogate for peripheral macrophages) (**C**), satellite glial cells and the subset of proliferating SGCs (**D**). Based on the Zeisel *et al* mouse DRG single cell RNA-seq dataset, several of these genes (*GAP43*, *NCAM1*, *NCAM2*, *APOE*, *FABP7*, *DBI*, *DHH*, *FBLN5*, *CEACAM10*) were found to be expressed in multiple cell types and not used for identifying putative increase / decrease of cell type proportions in culture. Neuronal and Schwann cell markers decrease markedly in culture, while fibroblast-like cell and SGC markers increased in culture. Macrophage / microglia markers profiled in this figure are not discriminative of solely that cell type in the DRG. NS: | SSMD | <= 2, NE: not systematically detected for that condition, N/A: not applicable because orthologous gene not identified in that species

**Figure 3.**
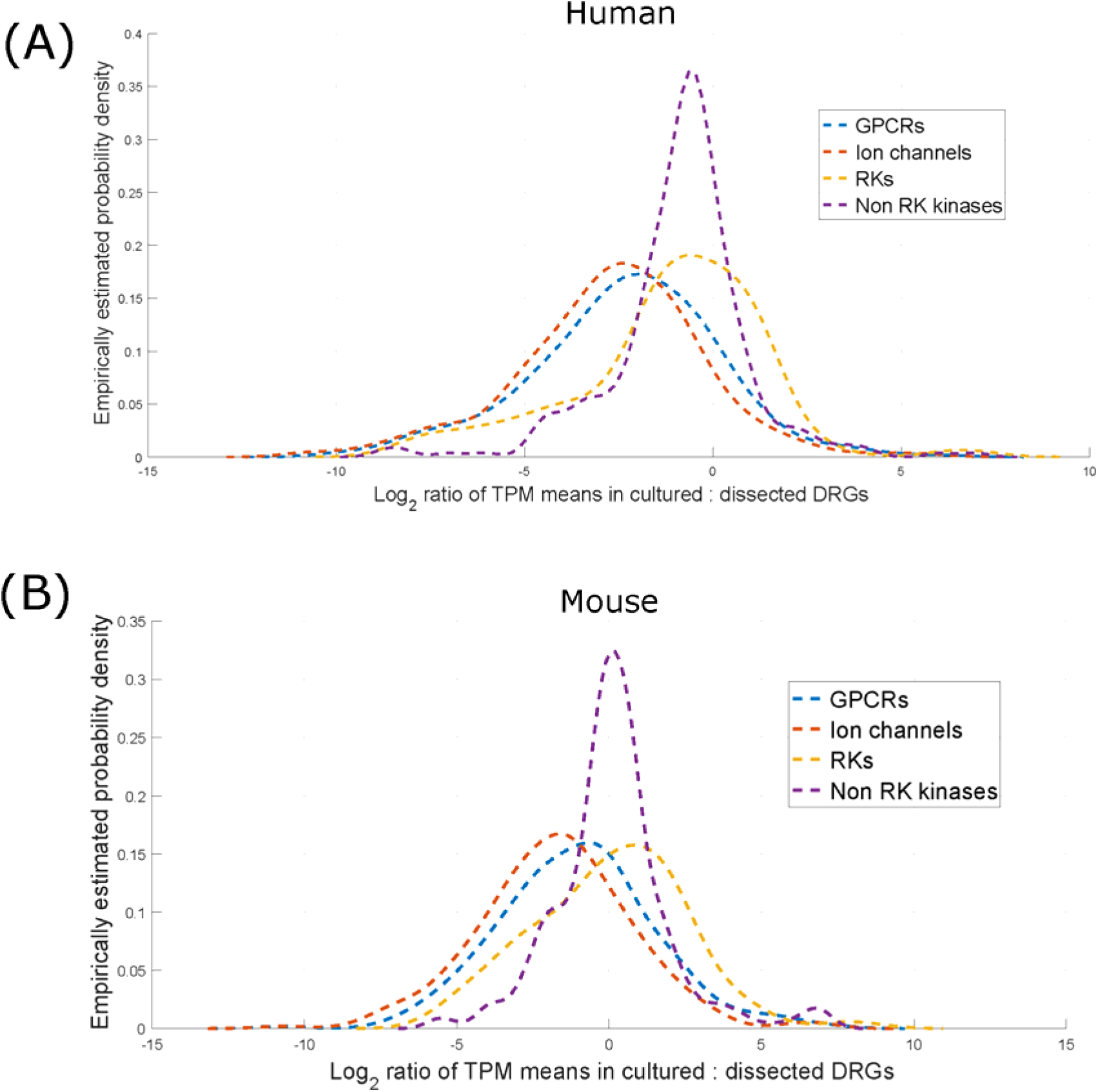
Empirical density distribution of log2 fold changes (ratio of means) for GPCRs, ion channels, RKs, and non-RK kinases in human (**A**) and mouse (**B**). RKs and kinases as a group are weakly de-enriched in human and weakly enriched in mouse cultures (in the context of mean expression). However, both GPCRs and ion channels are strongly de-enriched in both human and mouse cultures, likely because of the variety of these genes that are expressed in sensory neurons.

### Expression profiles of several pharmacologically relevant gene families show lower expression levels in DRG culture

A primary use of DRG cultures is to examine pharmacological effects of ligands for receptors with the assumption that this type of experiment reflects what occurs *in vivo* [35]. An underlying assumption of this type of experiment is that the presence or absence of a tested effect is reflected in consistent expression between *in vivo* and cultured conditions. To give insight into this assumption, we comprehensively cataloged expression of G-protein coupled receptors (GPCRs), ligand gated ion channels and receptor kinases (RKs) in native and cultured human and mouse DRG. To comprehensively characterize the changes in these gene families, we also characterized expression profiles of non-RK soluble kinases (**Supplementary file 1, Sheets 3-10**). We limited our soluble kinase comparisons to a well-characterized subset with clear mouse to human orthologs [32].

We find that a number of these genes are consistently detected in acutely dissected DRGs but not in culture. This was seen in human gene families of GPCRs (detected in intact DRG: 292; in culture: 190; out of which 176 were detected in both), ion channels (in intact DRG: 239, in culture: 179, in both: 172), RKs (in intact DRG: 68; in culture: 60; in both: 59), and non-RK kinases (in intact DRG: 286; in culture: 277; in both: 272); and a similar trend was observed in the mouse gene families as well. Since sensory neurons express a rich diversity of GPCRs and ion channels, the greater decrease in the number of systematically detected GPCRs and ion channels is likely the result of a proportional decrease of neurons in culture and/or decrease of gene expression in cultured neurons. Lists of systematically detected genes in these gene families are presented in **Supplementary File 1, Sheets 3-10**.

However, it is important to note that over 75% of the human genes in these families (human: 679 out of 885, mouse: 702 out of 824) that are consistently detected in intact DRG are still detectable in culture. This suggests that at single cell resolution, DRG cultures could be used as a surrogate for *in vivo* models in preclinical research for a majority of pharmacologically relevant molecular assays.

Next, for genes that are systematically detected in at least one condition, we identified the ones in these gene families that have | SSMD | > 2.0 (Tables 2 and 3, for human and mouse genes). Based on the SSMD values, while comparable numbers of GPCRs, ion channels and kinases were found to be decreased in cultured DRGs (GPCRs – human: 85, mouse: 95; ion channels – human: 109, mouse: 122; kinases – human: 106, mouse: 70), more mouse genes were detected to be systematically trending in the opposite direction as compared to their human counterparts (GPCRs – human: 7, mouse: 20; ion channels – human: 7, mouse: 14; kinases – human: 22, mouse: 66). As noted before, within-group variation is likely lower in mice due to controlled laboratory conditions and similar genetic backgrounds, and this enables us to detect more expression changes that have smaller effect sizes (as in the case of genes that are increased in cultured conditions).

**Table 2.**
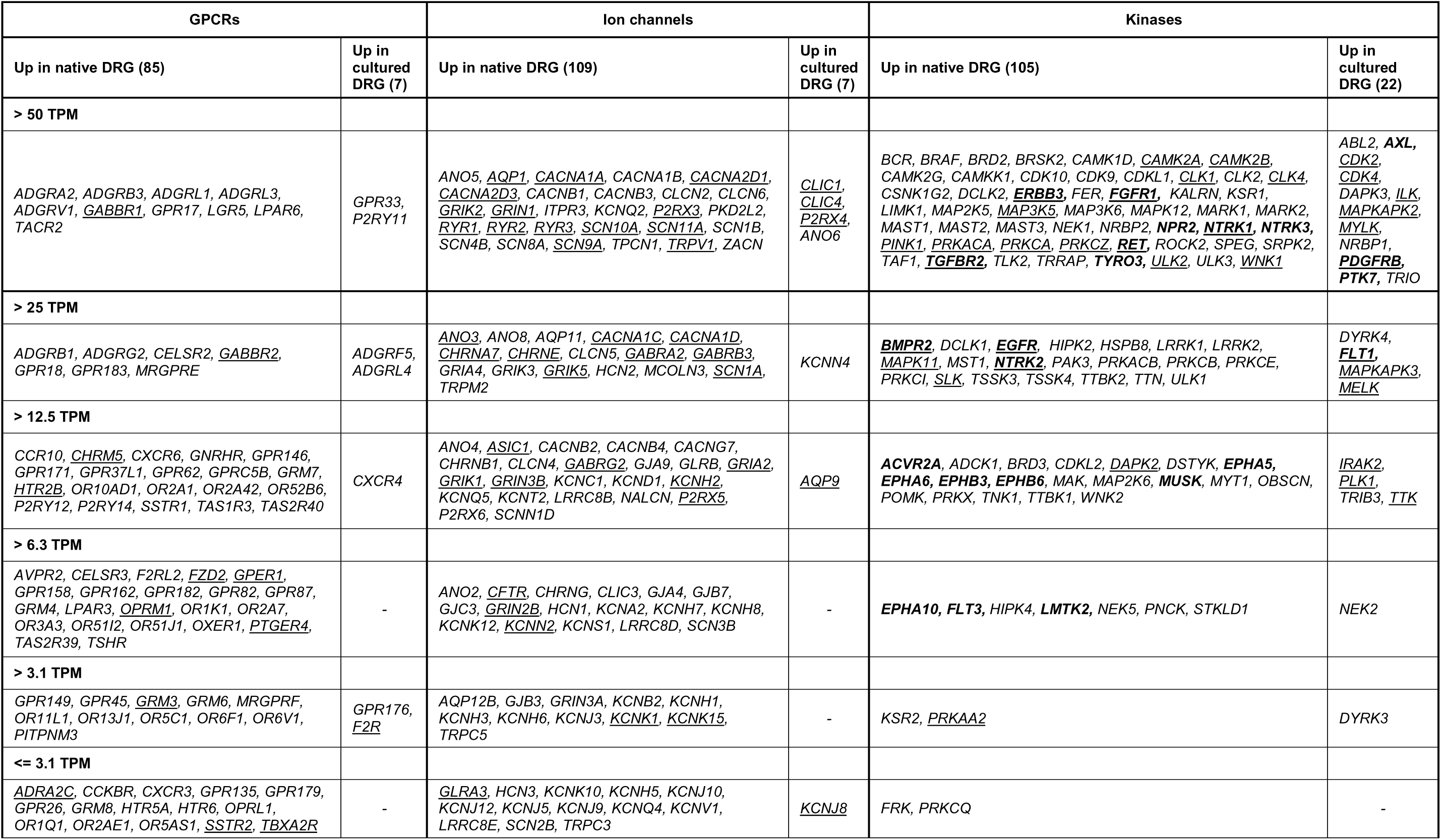
Differentially expressed pharmacologically relevant genes (RKs boldfaced) in intact vs cultured human DRGs with | SSMD | > 2, partitioned by mean TPM in condition of higher expression. The number of genes in each column is shown in parentheses. Genes known to be associated with pain from GWAS / functional association databases and the literature are underlined. Typically, smaller TPM brackets have higher technical variance.

**Table 3.**
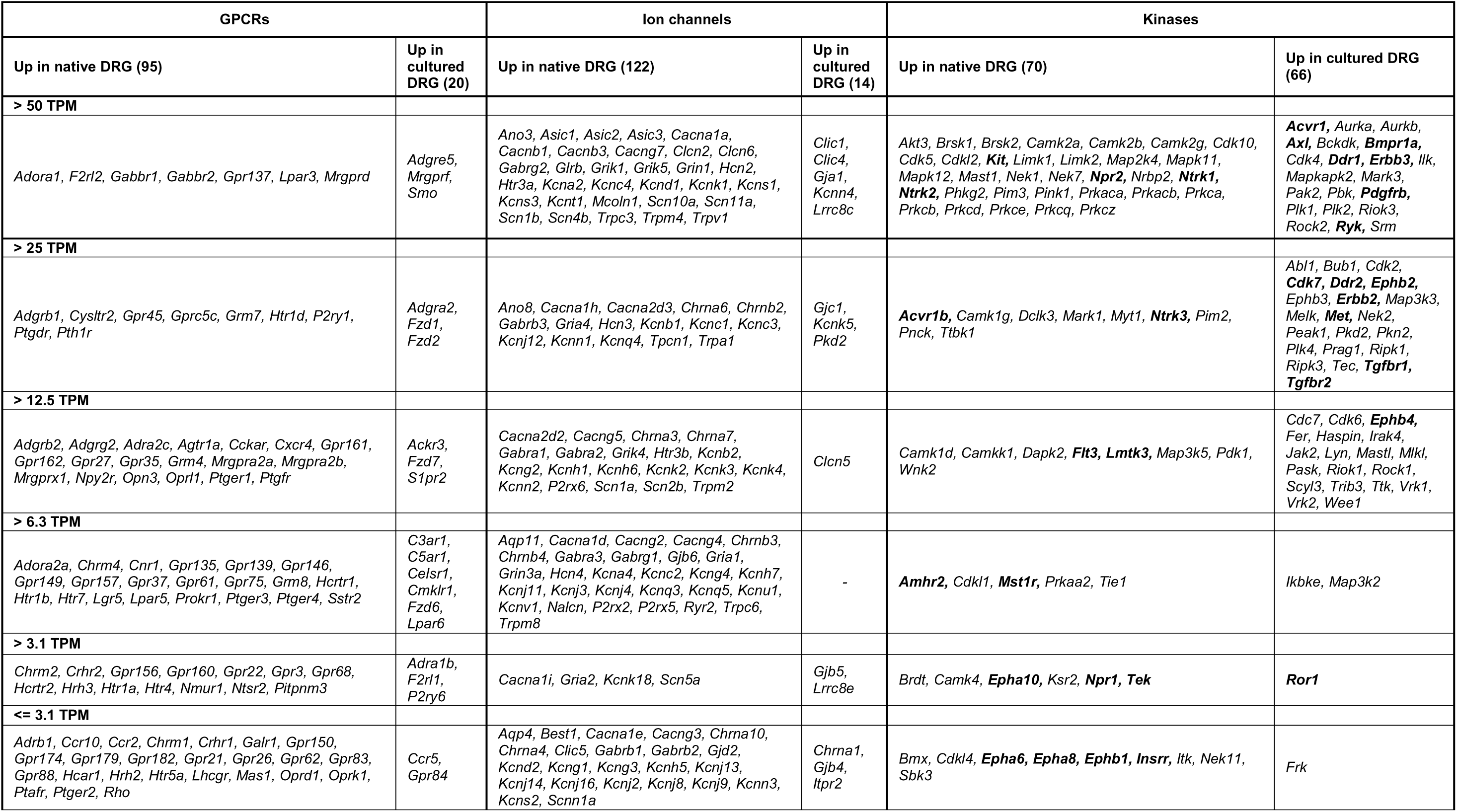
Differentially expressed pharmacologically relevant genes (RKs boldfaced) in intact vs cultured mouse DRGs, with | SSMD | > 2 partitioned by mean TPM in condition of higher expression. The number of genes in each column is shown in parentheses. Typically, smaller TPM brackets have higher technical variance.

We also characterized the degree of change in expression by estimating the probability density of the fold change (ratio of means) for all the genes in these families. The empirically estimated probability density for the ratio of means (intact DRG: cultured DRG) of the human and mouse pharmacologically relevant genes (Figure 3), shows a clear trend of decreased expression for a majority of the human ion channels and GPCRs.

Finally, we analyzed the trends in genes known to be involved in nociception, pain and neuronal plasticity. Genes with | SSMD | > 2.0 between conditions, and known to be associated with pain from the Human Pain Genetics Database, and the Pain – Gene association geneset (from the Comparative Toxicogenomics Database in Harmonizome [15; 49]), as well as from the literature, are underlined in Table 2. They identify pain-associated genes in these pharmacologically relevant families that change in expression between acutely dissected and cultured DRGs. Based on changes in consistent detectability between the two conditions, | SSMD | values > 2.0, or ratio of means > 2.0, changes in expression of several genes are discussed below.

#### Changes in human GPCRs

Several GPCRs involved in pro-inflammatory pathways, including *CCR1*, *CCRL2*, *CNR1*, *CXCR4*, *F2R*, *CHRM1* [66; 68] were found to increase in abundance in cultured DRGs. GPCRs found to be decreased in culture included *DRD5*, *HTR5A*, *HTR6*, and some metabotropic glutamate receptors (*GRM*s) like *GRM4, GRM5 (which was not detected) and GRM7*, all of which have been shown to be highly neural tissue enriched in humans (based on neural proportion score > 0.9 in Ray *et al* [47]). Their mouse orthologs have also been shown to be neuronally expressed in DRG single-cell RNA-seq experiments [58]. Many of these and other GPCRs changing in abundance between acutely dissected and cultured DRGs (Table 2, **and Supplementary File 1 Sheet 3**) have been noted as potential targets for pain treatment [7; 14; 34; 52]. Therefore, our findings suggest that under certain culture conditions false negatives could arise for these targets.

#### Changes in mouse GPCRs

Pro-inflammatory mouse GPCRs were also found to be increased in cultured DRGs, including *Ccr5*, *Cxcr6*, *F2r*, and *F2rl1* [29; 56; 60]. Several neuronally expressed mouse GPCRs (based on Usoskin et al [58]), including *Chrm2*, *Htr1a*, *Htr2c*, *Htr7*, and metabotropic glutamate receptors like *Grm4* were found showed higher expression in acutely dissected DRGs (Table 3, **and Supplementary File 1 Sheet 4**). Many of these genes in the human and mice datasets were from orthologous families of receptors, including cytokine receptors, the protease activated receptor (PAR) family (*F2R*), 5-HT receptors, and metabotropic glutamate receptors.

#### Changes in human ion channels

Among the ion channels increased in abundance in cultured DRGs were the chloride intracellular channels *CLIC1* and *CLIC4*, gap junction protein *GJA1*, *KCNG1* (K_v_6.1), *KCNJ8* (K_IR_6.1), *KCNN4* (K_Ca_4.2), and *P2RX4*, *TRPV4*, and voltage dependent anion channels *VDAC1* and *VDAC2*. Interestingly, many of these ion channels are involved in membrane potential hyperpolarization, suggesting a potential compensatory mechanism to suppress excitability. Neuronally expressed voltage gated calcium channels such as *CACNA1B*, *CACNA1F*, *CACNA1I*, *CACNAG5*, *CACNAG7* and *CACNAG8*; glutamate ionotropic receptors *GRIA2* and *GRIN1*; voltage gated potassium channels *KCNA1*, *KCNA2*, *KCNB2*, *KCNC3*, *KCND1*, *KCND2*, *KCNH2*, *KCNH3*, *KCNH5*, *KCNJ12*, *KCNK18*, *KCNQ2*, *KCNT1*, *KCNV1*; purinergic receptors *P2RX2* and *P2RX5*; and voltage-gated sodium channels *SCN1A*, *SCN4A*, *SCNN1A* and *SCNN1D* were found to be increased in intact DRGs. (Table 2, **and Supplementary File 1 Sheet 5**)

#### Changes in mouse ion channels

Changes in mouse ion channel genes were also quantified. (Table 3, **and Supplementary File 1 Sheet 6**). Genes increased in DRG cultures included several of the same families seen in human, such as chloride intracellular channels *Clic1* and *Clic4*; gap junction proteins *Gja1*, *Gja3*, *Gjb3*, *Gjb4*, *Gjb5* and *Gjc1*; the glutamate ionotropic receptor *Grik3*; voltage-gated potassium channels *Kcnk5* (K_2p_5.1) and *Kcnn4* (K_Ca_4.2); and purinergic receptors *P2rx1*, *P2rx7*. Among ion channels decreased in culture were the chloride intracellular channel *Clic3* and *Clic5*; voltage-gated calcium channels *Cacna1i*, *Cacna1s*, *Cacng3*; cholinergic receptors *Chrna10*, *Chrna6*, *Chrnb3* and *Chrnb4*; glutamate ionotropic receptors *Grik1*, *Grin2c*; 5-HT receptors *Htr3a* and *Htr3b*; voltage-gated potassium channels *Kcnd2*, *Kcng3*, *Kcng4*, *Kcnj11*, *Kcnj13*, *Kcnn1*, *Kcnn2* and *Kcns1*; *P2rx2*; voltage-gated sodium channels *Scn1A* and *Scn11A*; and TRP channels *Trpm2* and *Trpm8*. Most of these genes are well known to be neuronal in expression [58]. Overall, the ion channel subfamilies changing in expression in culture in both species were similar and included primarily voltage-gated calcium/potassium/sodium channels, purinergic receptors and gap junction proteins.

#### Changes in human RKs and other kinases

We found that the neuronally expressed genes from the *NTRK* family (*NTRK1, NTRK2, NTRK3*) and the CAMK family (*CAMK1D*, *CAMK1G*, *CAMK2A*, *CAMK2B*, *CAMK2G*, and *CAMKK1*) were decreased in culture in the human DRG. (Table 2, **and Supplementary File 1 Sheets 7 and 9**)

#### Changes in mouse RKs and other kinases

Consistent with what we found in the human cultures, we identify decrease in abundance in neuronally expressed Ntrk family (*Ntrk1*, *Ntrk2*, *Ntrk3*) and Camk (*Camk1g*, *Camk2a*, *Camk2b*) family genes. The changes in the Ntrk family, responsible for neurotrophin signaling in adult DRG neurons, demonstrates a consistent inter-species trend in culture. Consistent trends in the Camk family genes, which play a vital role in Ca^2+^-dependent plasticity in the brain [12] and in nociceptors [9; 10; 20], also show conserved patterns in the DRG cultures. (Table 3, **and Supplementary File 1 Sheets 8 and 10**)

### Neuronal injury and inflammation markers were increased in human and mouse DRG cultures

Dissection of the DRG causes an axotomy that may induce cells to take an inflammatory phenotype as is seen *in vivo* after peripheral nerve injury [38]. However, this idea has not been systematically investigated on a genome-wide scale. As shown in Figure 4A-C, many genes associated with inflammation and cell proliferation, neuronal injury and repair, and immune signaling and response, including cytokines [1; 8; 61] and matrix metalloproteases [28] associated with neuropathic pain, were differentially expressed in human and mouse DRG cultures with respect to intact DRG. As extremal examples of gene expression changes in our datasets, *IL6* and *MMP9* mRNA expression were increased 100 fold or more in human DRG culture (Fig 4C).

**Figure 4.**
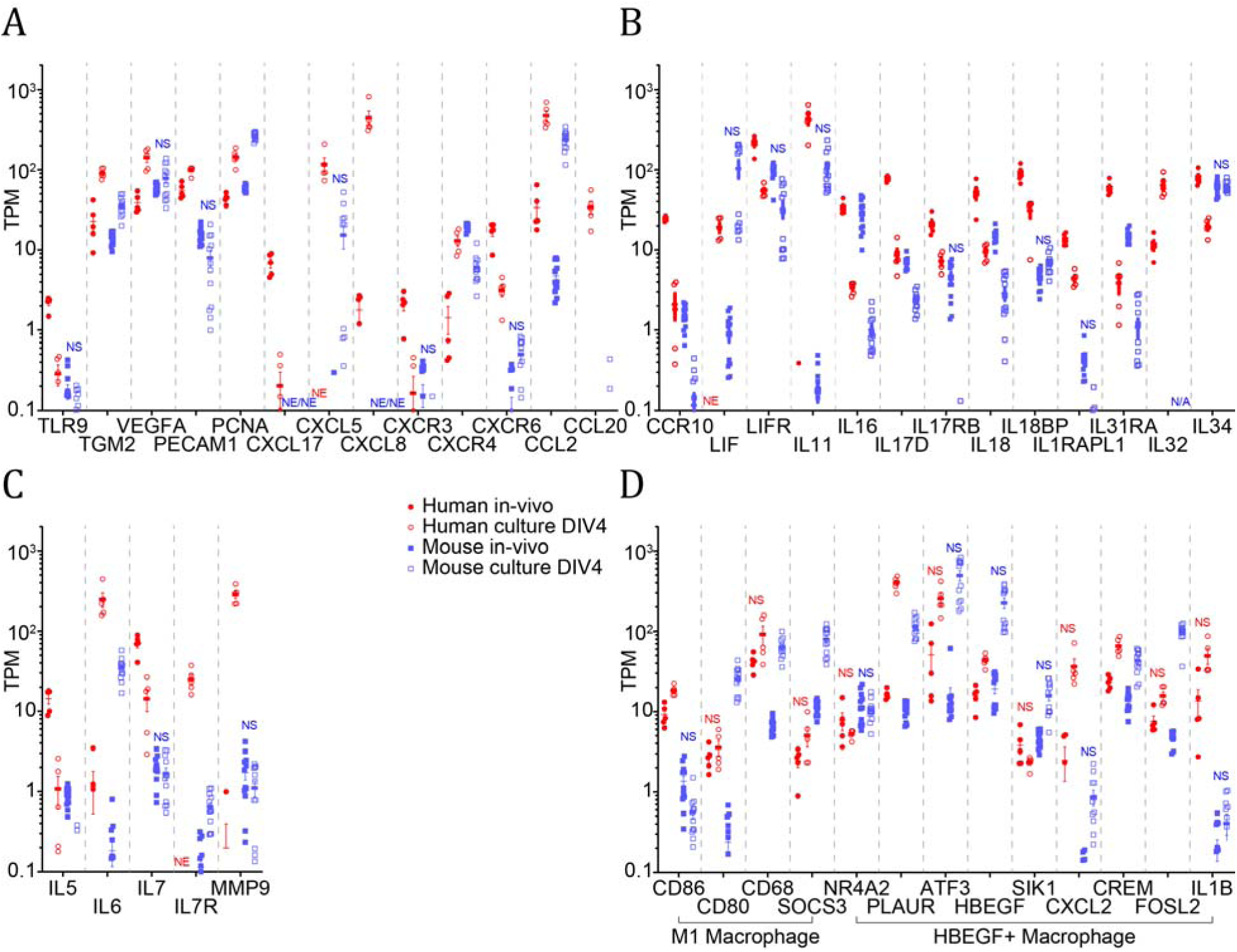
Expression levels in human and mouse intact vs. cultured DRGs. A wide diversity of genes involved in inflammation and proliferation, nerve and neuronal injury and repair, and immune signalling and response are profiled (**A, B, and C)**. Key expressed genes for M1 macrophages and *HBEGF*+ macrophages are also shown (**D**). NS: | SSMD | <= 2, NE: not systematically detected for that condition, N/A: not applicable because orthologous gene not identified in that species.

Since several of these genes are increased or decreased in cultured samples, we used the mouse DRG single cell RNA-seq profiles [58] to putatively identify cell types of expression among cells constituting the DRG (**Supplementary File 1 Sheet 11**). Indeed, we find that genes primarily expressed in neurons and Schwann cells decrease in abundance, even if they are involved in pro-inflammatory signaling, since it is likely that these cells types are reduced in frequency in DRG cultures. Interestingly, several genes predicted to be primarily expressed in immune cells (*TLR9*, *CXCR3*) and in vascular cells (*IL18BP*, *CXCL17*) were found to be reduced in relative abundance in cultures, suggesting that potential increase in immune and vascular cell proportions in culture are limited to certain cell subtypes in these categories.

Multiple subtypes of macrophages are involved in inflammatory processes and can be identified with specific markers [23]. In human and mouse, key M1 macrophage genes *CD68, CD80, and SOCS3* were all upregulated in culture compared to intact ganglia. As identified in a recent study, HBEGF+ inflammatory macrophages are responsible for fibroblast invasiveness in rheumatoid arthritis patients [30]. We noted that multiple genes expressed in this specific subtype of macrophage (*PLAUR*, *HBEGF*, *CREM*) were increased in human and mouse DRG cultures, suggesting that this particular subtype of macrophage may be present in DRG cultures from both species (Fig 4D).

While specifically identifying the exact subtype of immune cell involved is outside the scope of our bulk RNA-sequencing assay, our findings reveal clearly that many genes involved in neuronal injury, cell proliferation and inflammation, and immune signaling and response are increased in DRG cultures.

### Similarities and differences between human and mouse DRG culture transcriptomes in the context of intact DRG transcriptomes

Complicated orthologies and differential evolutionary dynamics between human and mouse gene families [67], and gaps in human to mouse orthology annotation [37] make comparative transcriptomic comparisons difficult between human and mouse transcriptomes. We have previously made similar comparisons between native human and mouse acutely dissected DRGs [47], finding overall similarities, but also some changes in gene expression. Since we are analyzing changes in expression at the level of individual genes (such as pharmacologically relevant ones), we limited our analysis to changes in expression in GPCRs, ion channels, and kinases in DRG cultures for tractability.

We calculated trend scores for each GPCR, ion channel, RK, and non-RK kinase, after eliminating genes from the analysis with complicated orthologies between humans and mouse (**Supplementary File 1, Sheets 12-15**). We find a weak correlation between trend scores of human genes and their mouse orthologs in GPCRs (Pearson’s R: 0.19, one tailed test *p* value: 0.0008) and ion channels (Pearson’s R: 0.15, one tailed test *p* value: 0.012). For specific genes, we find consistent increase (eg. *F2R*, *GPRC5A*, *TRPV4*) or decrease (eg. SCN subfamily members, GABAR subfamily members) in cultured samples for both species. This suggests that expression patterns across cell types, which potentially contributes to the trend scores, is likely conserved in these genes across species. However, in several cases, genes may not be consistently detectable in one species but present in one or more conditions in the other (eg. *CHRM5* only detectable in human DRGs, *CHRNB4* only detectable in mouse DRGs). *TRPC5* is expressed at low levels in 3 intact mouse DRG samples, but significantly increased in expression in all human samples. Additionally, several genes are expressed in both species, but have opposing expression trends across intact and cultured DRGs (eg. *ACKR4* and *CXCR6* decreased in human cultures, but increased in mouse cultures). Such changes are likely due to evolutionary divergence between species in gene expression across cell types, and/or differential transcriptional regulation between species. Both of these involve regulatory evolution.

**Supplementary File 1 Sheets 12 – 15** profile members of these gene families, their trend scores, and the number of human and mouse samples where they are detectable. This provides a roadmap for identifying genes changing in cultured versus intact DRGs across species; and creates a resource for the neuroscience community interested in performing molecular assays in cultured DRGs on these genes.

Since several members of the MRGPR family do not have a one-to-one orthology between human and mouse genes, they were not included in the trend score calculation tables (**Supplementary File 1 Sheet 12**). TPM values from human and mouse cultures for all members of this gene family are presented in their own table because this family of genes plays an important role in sensory neuroscience (Table 4).

**Table 4.** MRGPR/Mrgpr family gene expression levels in human and mouse

| Human |  |  | Mouse |  |  |
| --- | --- | --- | --- | --- | --- |
| Gene name | Mean TPM in intact DRGs | Mean TPM in cultured DRGs | Gene name | Mean TPM in intact DRGs | Mean TPM in cultured DRGs |
| *MAS1* | 0.02 | 0.06 | *Mas1* | 0.75 | 0.05 |
| *MAS1L* | 0.0 | 0.0 | No *MAS1L* ortholog | - | - |
| *MRGPRD* | 0.85 | 0.21 | *Mrgpra1* | 0.77 | 0.04 |
| *MRGPRE* | 31.41 | 1.53 | *Mrgpra2a* | 20.91 | 0.67 |
| *MRGPRF* | 5.45 | 1.84 | *Mrgpra2b* | 23.85 | 0.96 |
| *MRGPRG* | 0.00 | 0.00 | *Mrgpra3* | 24.50 | 0.02 |
| *MRGPRX1* | 4.45 | 0.48 | *Mrgpra4* | 0.48 | 0.01 |
| *MRGPRX2* | 0.00 | 0.00 | *Mrgpra6* | 0.00 | 0.00 |
| *MRGPRX3* | 0.32 | 0.08 | *Mrgpra9* | 0.98 | 0.01 |
| *MRGPRX4* | 0.12 | 0.00 | *Mrgprb1* | 0.15 | 0.03 |
|  |  |  | *Mrgprb2* | 0.09 | 0.00 |
|  |  |  | *Mrgprb3* | 0.00 | 0.00 |
|  |  |  | *Mrgprb4* | 7.33 | 0.00 |
|  |  |  | *Mrgprb5* | 9.14 | 0.04 |
|  |  |  | *Mrgprb8* | 0.02 | 0.00 |
|  |  |  | *Mrgprd* | 74.20 | 0.03 |
|  |  |  | *Mrgpre* | 21.22 | 19.68 |
|  |  |  | *Mrgprf* | 7.91 | 73.72 |
|  |  |  | *Mrgprg* | 0.00 | 0.00 |
|  |  |  | *Mrgprh* | 0.23 | 0.04 |
|  |  |  | *Mrgprx1* | 18.86 | 0.82 |
|  |  |  | *Mrgprx2* | 0.04 | 0.01 |

### Similarity and differences between cultured DRG transcriptomes across different labs

For mouse, experiments were performed in 2 labs (Gereau lab – sample ids with a “g” suffix; and Price lab – sample ids with a “p” suffix, Figure 1B) independently. Although both labs used the same strain of mouse, both intact and cultured DRGs had a clear transcriptome difference between the two labs. This is likely caused by environmental differences between animal facilities. Additionally, while changes in gene expression levels are well known to be different across inbred mouse strains [57], recent research suggests that even for inbred mouse strains separated for over hundreds of generations, mutation profiles diverge and can cause different outcomes in molecular assays, and have been shown to cause changes in immune function related genes [11]. Surprisingly, we saw that inter-laboratory transcriptome differences in cultured mouse DRGs were smaller in cultured samples with respect to acutely dissected DRGs despite differences in culturing protocols (e.g. without nerve growth factor (NGF) in the Gereau lab, and with NGF in the Price lab). This is likely due to the fact that neurons have the most plastic molecular profiles, and putatively decline in proportion in cultured DRGs.

The small amount of changes in DRG culture between the two laboratories can be summarized as follows. A large amount of overlap was found in systematically detected genes for GPCRs (consistently detected in Price lab culture: 191, in Gereau lab culture: 214, in both labs’ cultures: 183; Figure 5), ion channels (consistently detected in Price lab culture: 204, in Gereau lab culture: 217, in both labs’ cultures: 200; Figure 6), and RKs (consistently detected in Price lab culture: 58, consistently detected in Gereau lab culture: 65, consistently detected in both labs’ cultures: 57; Figure 7). Among the genes that have a greater than 2-fold change in expression is *Ngfr* (mean TPM in Price lab: 973, in Gereau lab: 438), potentially due to the use of NGF in the culturing process in the Price lab. Several genes that were detected in one or both labs’ cultures had laboratory-specific expression changes with | SSMD | > 2, and are noted in Figures 5, 6 and 7.

**Figure 5.**
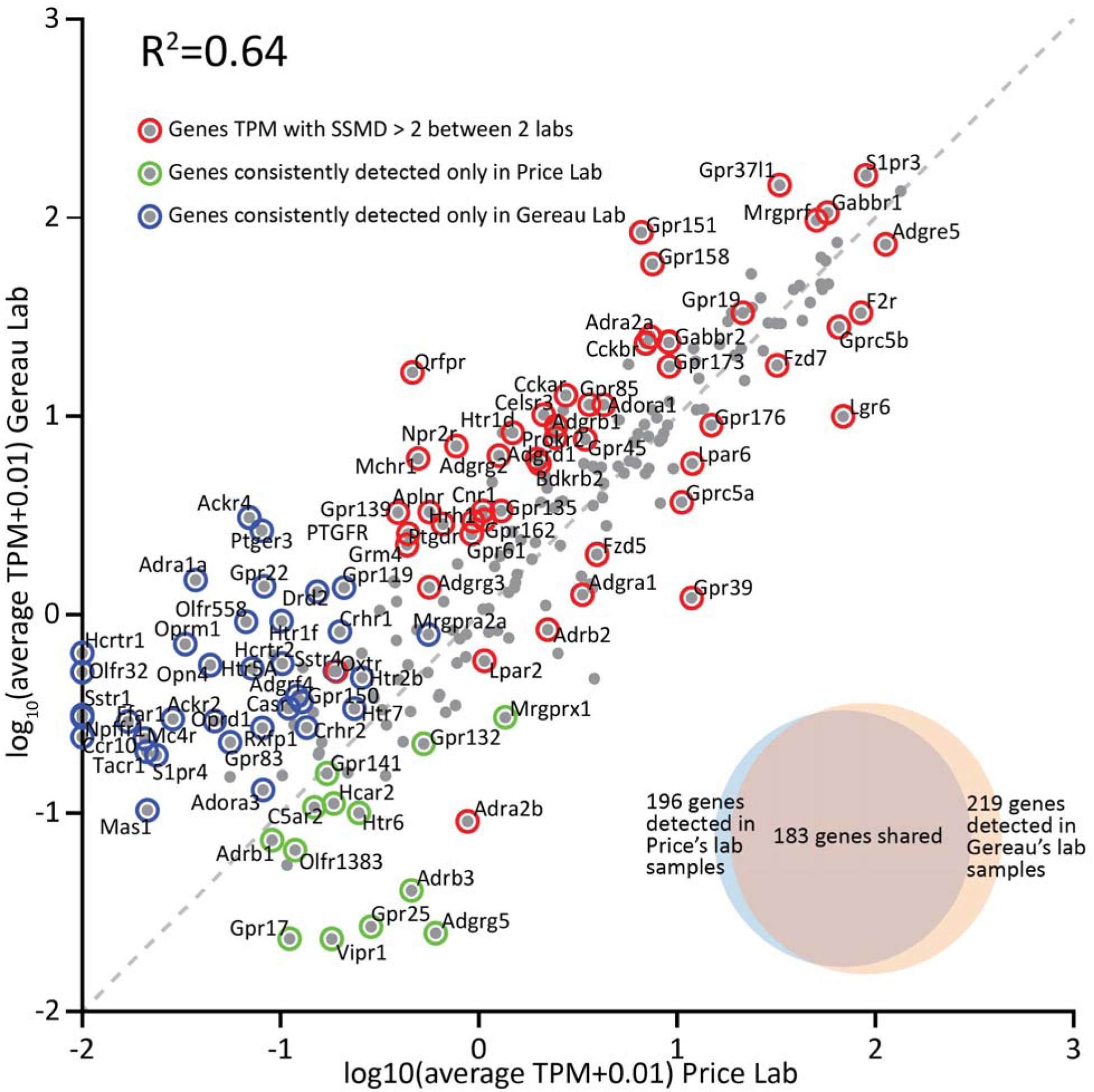
Scatter plot and Venn diagrams showing differential expression of GPCR genes in culture between the Price and Gereau lab. GPCR gene expression levels from each lab were plotted on the X and Y axes showing moderate correlation of expression profiles between labs (Pearson’s R squared: 0.64, p << 0.01).

**Figure 6.**
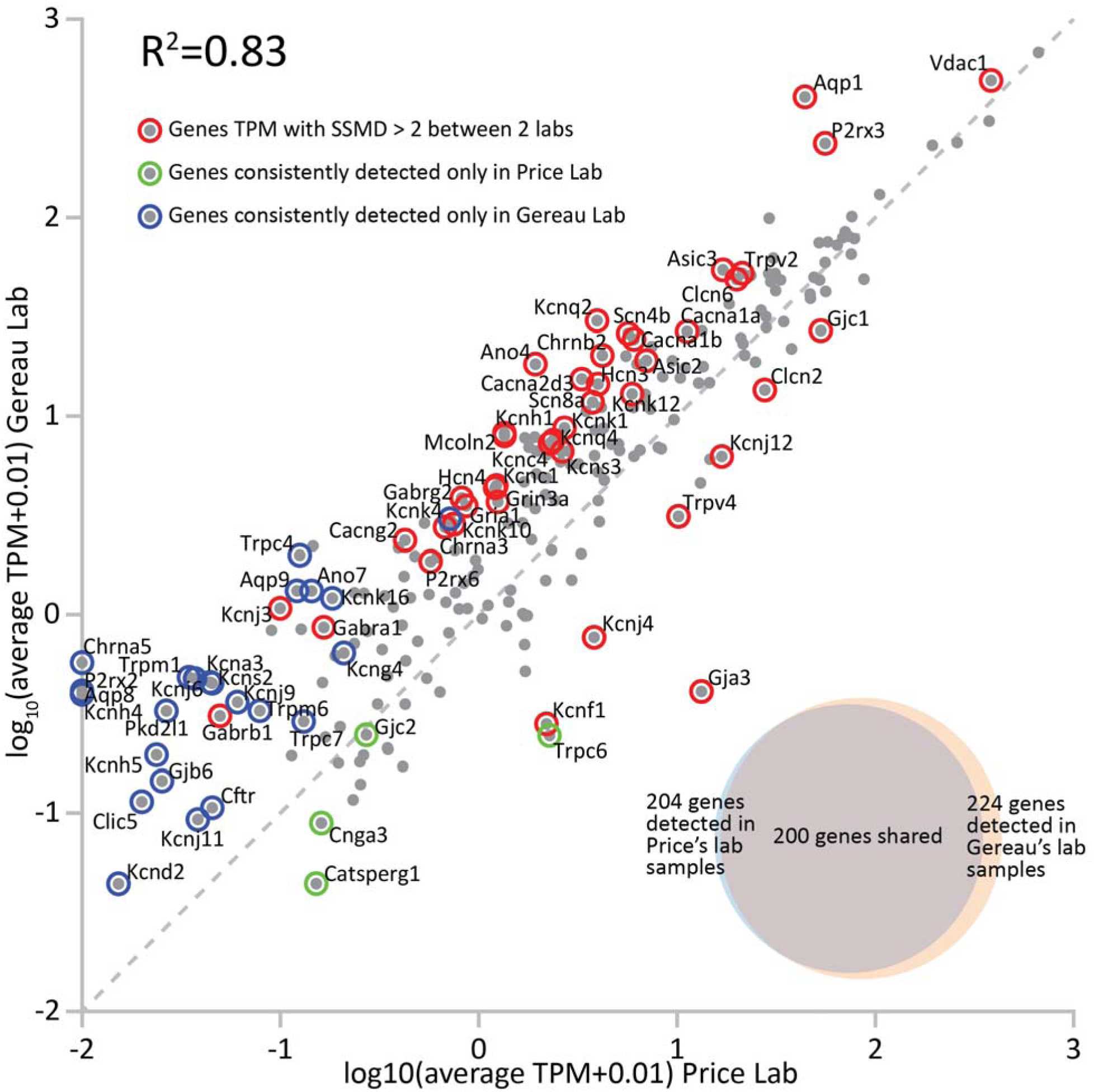
Scatter plot and Venn diagrams showing differential expression of ion channel genes in culture between the Price and Gereau lab. Ion channel gene expression levels from each lab were plotted on the X and Y axes showing strong correlation of expression profiles between labs (Pearson’s R squared: 0.83, p << 0.01).

**Figure 7.**
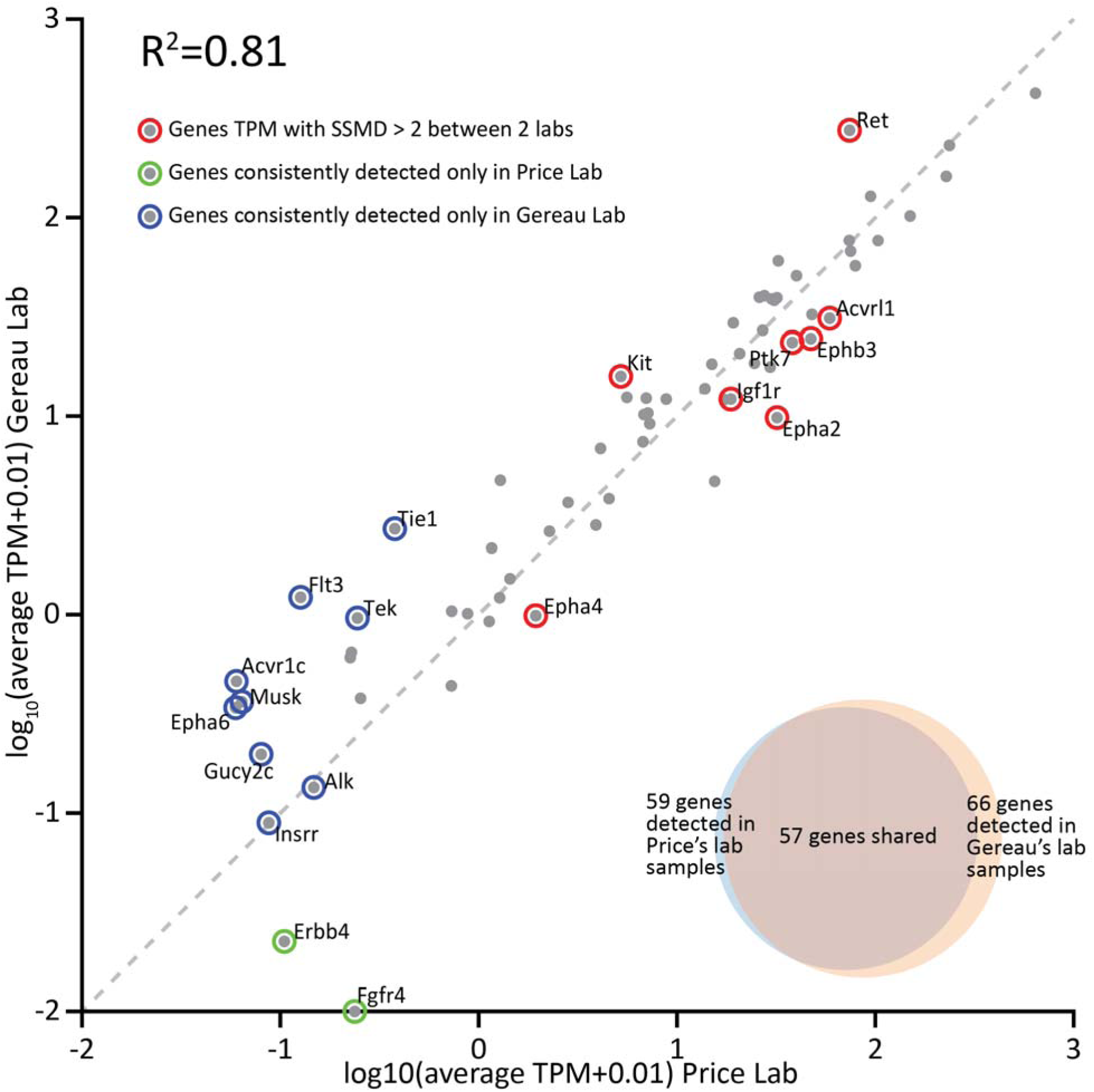
Scatter plot and Venn diagrams showing differential expression of RK genes in culture between the Price and Gereau lab. RK gene expression levels from each lab were plotted on the X and Y axes showing strong correlation of expression profiles between labs (Pearson’s R squared: 0.81, p << 0.01). Alk and Insrr are plotted on the diagonal, but marked as consistently detected only in Gereau lab samples. This is because they have comparable mean TPMs in samples from both labs, but are only consistently detected (in 5 or more samples out of 6) in the Gereau lab.

## Discussion

While DRG cultures are used for electrophysiology, Ca^2+^ imaging, cell signaling and a variety of other types of physiological studies, we are unaware of any previous studies that have used genome-wide technologies to characterize transcriptomes between acutely dissected and cultured DRGs. While in vivo cross-species comparisons have previously been made [47], *in vitro* transcriptome comparisons between mice and human DRGs have not been performed, despite the obvious need for such knowledge given the reliance on the mouse model for both target and drug discovery work in the pain area [14; 35; 46; 59; 70]. Certain perturbation studies [43] like gene expression knockdowns, DNA editing or optogenetic optimization [36], especially in the context of human research, cannot be performed *in vivo*, causing DRG cultures to be essential to human clinical translational research. Our work, therefore, gives fundamental new insight into some of the most commonly used model systems in the pain field with important implications for future work.

We reach two major conclusions based on the experiments we have presented here. First, while many pharmacologically meaningful features of the DRG are well-conserved from mouse to human, there are some important differences that need to be considered in future experimental design. Moreover, there are a small but potentially important number of human receptors that simply cannot be studied in culture systems that may be good targets for drug discovery.

Second, mouse and human DRG cultures take on an inflammatory-like transcriptomic phenotype that shares some qualities with transcriptomic changes that occur during the pathogenesis of neuropathic pain [1; 28; 31; 39]. Therefore, the cultured DRG system may reflect certain clinical features that would be advantageous for neuropathic pain mechanism and/or drug discovery, especially in humans where these samples are not readily available except under very unique circumstances [39].

A potential critique of using primary neuronal cultures to test pharmacological targets is that cultures are not an accurate representation of native tissue. While there were some specific genes that did not appear in culture when compared to native tissue and vice versa, the main difference between the two conditions was in the expression level of each gene, which our data strongly suggests is due to the change in the proportion of cell types. Specifically, the proportion of neurons in culture was decreased when compared to macrophage, fibroblast and SGCs. To this end, when specific pharmacological targets are being tested in either mouse or human cultures it is important to check that these targets remain expressed and our work provides a comprehensive resource to do this in both species (**Supplementary File 1**). Because pharmacology is the most common use of cultured DRG, we focused our analysis on pharmacologically relevant targets. Interestingly, both mouse and human cultures displayed an increase in M1 and HBEGF+ macrophage [30] markers when compared to native tissue. This change suggests an increase in the inflammatory macrophage population in culture. We predict that this shift is due to phenotypic shifts of tissue resident macrophages caused during the dissociation and culturing process, potentially replicating a nerve injury phenotype [1; 8; 27; 28; 61]. The presence of these cell types in culture could be employed to further study how macrophages and sensory neurons interact.

When comparing the gene expression differences between mouse and human cultures we found some differences, consistent with our previous analysis of native DRG species differences [47]. Families of genes remained consistently expressed in both species following dissociation and culturing protocols, but individual genes of the same family varied in whether they were present in either mouse or human. For example, *Kcna1* was systematically detected only in mouse DRG cultures. Therefore, while most ion channel types are likely to be equally represented in both human and mouse DRG neurons, there is a substantial chance that the specific subtypes of channels that make up those conductances will be different between species, and such changes may be present both *in vivo* and in culture. In fact, studies focusing on exactly this question for voltage gated sodium channels in DRG between rat and human have found qualitative similarities but key differences that are almost certainly due to differences in expression between species [70]. This is a critical distinction for pharmacology because a primary goal in therapeutic development is ion channel subtype specific targeting [46]. Our findings demonstrate that it is vital to understand these similarities and differences when choosing a model system to study a particular target and, critically, we provide a comprehensive resource to do this. From a discovery perspective, studies performed *in vitro* in mouse neuronal cultures likely remain a valid and reliable option for researchers as the *families* of ion channels, GPCRs, and RKs are well conserved from mice to humans (**Supplementary File 1, sheets 12-15**).

Our study has several limitations to acknowledge. The first is the choice of time point for the cultured DRG RNA-seq studies. We chose 4 DIV for our studies. Given the literature on biochemical and Ca^2+^ imaging studies (which is too extensive to cite) we think that our findings will provide a substantial resource for studies of this nature as most of them are done between 3 and 7 DIV. This can also be said for many electrophysiological studies on human DRG neurons as most investigators do experiments on these neurons over many days, with 4 DIV falling in the middle of the experimental spectrum for this small, but growing, body of work. The exception is mouse DRG electrophysiology where the vast majority of this very large literature has been done at 24 hrs after culturing. It is possible that some of the changes we observe at 4 DIV are not present at less than 1 DIV and/or that other differences are observed at this early time point. Another limitation is that changes in mRNA expression in culture may not represent differences in functional protein because some of these proteins may have long half-lives. In such a scenario, a down-regulation of mRNA would not lead to any difference in functional protein over the time course of our experiment (4 DIV). This can only be addressed with proteomic or physiological [50; 70] methods, which we have not done. Finally, we have relied on bulk RNA sequencing in the work described here. We acknowledge that single cell sequencing would yield additional insights that will be useful for the field. This will be a goal of future work.

We have focused on using DRGs from uninjured mice and from human organ donors without a history of chronic pain. Many studies have demonstrated that cultured DRG neurons from mice with neuropathic pain retain some neuropathic qualities *in vitro*, in particular spontaneous activity in a sub-population of nociceptors [2; 33; 39; 63; 65]. This also occurs in human DRG neurons taken from people with neuropathic pain [39]. Our transcriptomic studies suggest that cultured DRG neurons from normal mice and human organ donors show some transcriptomic changes consistent with a neuropathic phenotype, but these neurons likely do not generate spontaneous activity. Some transcriptomic changes were found in mice but not conserved in humans. An example is caspase 6 (*Casp6*) which has been implicated in many neuropathic pain models in mice [4; 5; 33]. This gene was over 3 fold increased in mouse DRG culture but unchanged in human DRG cultures.

An interesting observation emerging from our work is that some macrophages are apparently present in DRG cultures and this macrophage phenotype is inflammatory in nature. An emerging literature describes DRG resident macrophages as key players in development of many chronic pain states, including neuropathic pain [23; 27; 39; 51; 61]. In future studies it may be possible to manipulate these macrophages to interact with DRG neurons in culture to push this neuropathic pain phenotype further toward the generation of spontaneous activity. Such studies could allow for the generation of a neuropathic pain model *in vitro*. Such an advance would be particularly useful for the human organ donor DRG model, especially considering that neuropathic pain in patients is associated with a macrophage transcriptomic signature, at least in males [39].

We have comprehensively characterized transcriptomic changes between native and cultured mouse and human DRG. Our overarching conclusion is that these tissues are similar between the two species, suggesting that discovery work that is largely done in mice faithfully models many physiological characteristics of human DRG neurons. However, there are important differences between species and between native and cultured conditions, with minimal impact of the type of culturing protocol used. Our resource brings these differences to light allowing for appropriate model system choice and delineation of pharmacological divergences.

